# Persistent alterations in plasma lipid profiles prior to introduction of gluten in the diet associate with progression to celiac disease

**DOI:** 10.1101/419416

**Authors:** Partho Sen, Cecilia Carlsson, Suvi M. Virtanen, Satu Simell, Heikki Hyöty, Jorma Ilonen, Jorma Toppari, Riitta Veijola, Tuulia Hyötyläinen, Mikael Knip, Matej Orešič

## Abstract

**Background and Aims:** Celiac disease (CD) is a chronic enteropathy characterized by an autoimmune reaction in the small intestine in genetically-susceptible individuals. Gluten is the required environmental trigger of clinical CD, but the underlying causes of the autoimmune reaction remain unknown. Herein, we apply lipidomics to elucidate the early events preceding clinical CD in a prospective study of children observed from birth until diagnosis of CD and subsequent introduction of a gluten-free diet.

**Methods:** Mass spectrometry–based lipidomics profiling was applied to a longitudinal series of 233 plasma samples from the Type 1 Diabetes Prediction and Prevention (DIPP) study, spanning the period between birth and the introduction of a gluten–free diet following CD diagnosis (n=23 CD progressors, n=23 controls matched for gender, HLA risk, period of birth, and age).

**Results:** 23 children progressed to CD at a mean age of 4.8 years. They showed increased amounts of triacylglycerols (TGs) of low carbon number and double bond count and a decreased level of phosphatidylcholines by 3 months of age as compared to controls. These differences were exacerbated with age but were not observed at birth. No significant differences were observed in essential (dietary) TGs such as those containing polyunsaturated fatty acids.

**Conclusion:** Our findings suggest that abnormal lipid metabolism associated with development of clinical CD may occur prior to the introduction of gluten to the diet. Moreover, our data suggest that the specific TGs found elevated in CD progressors may be due to a host response to compromised intake of essential lipids in the small intestine, requiring *de novo* lipogenesis.

## INTRODUCTION

Celiac disease (CD) is a chronic, systemic, autoimmune enteropathy triggered by dietary gluten and related prolamines from rye and barley in genetically-susceptible individuals^1-6^. About 90-95% of CD patients express HLA-DQ2 protein, while the remaining (5-10%) express HLA-DQ8^7, 8^. CD is characterized by a wide range of gastro-and extraintestinal symptoms that include diarrhea, weight loss, abdominal distention, malabsorption and iron-deficiency anemia^1, 4, 9^. Serologic tests such as measurement of serum IgA and/or IgG tissue transglutaminase antibodies, IgA endomysial antibodies, deamidated gliadin peptide antibodies (IgG class) are performed for screening and diagnosis of CD. In addition, a biopsy of small intestine is still required in many countries to confirm the diagnosis^2, 4^.

The incidence of CD and other autoimmune diseases such as type 1 diabetes (T1D) have been increasing in children and adults over the past decades^10, 11^. The occurrence of these autoimmune diseases is higher in the Nordic countries^12^ than elsewhere, with the highest prevalence of CD occurring in Sweden (29/1000 by the age of 12) and the highest incidence rate of T1D occurs in Finland (64/100000/year for children under 15 years of age)^13, 14^. Approximately 10% of patients with T1D develop overt CD^15, 16^. On the other hand, people with CD are at-risk for T1D before 20 years of age^17^. These autoimmune diseases share common, predisposing alleles in the class II HLA-region as the DR3-DQ2 and DR4-DQ8 haplotypes^18-21^.

Recent studies show that, in addition to genetic predisposition and exposure to dietary gluten, other factors such as the composition of the intestinal microbiota, birth delivery mode, infant feeding and the use of antibiotics may also affect the onset of CD^1, 22^. Thus, the early pathogenesis of CD is still poorly understood^4, 23, 24^ and the identification of molecular signatures associated with progression to overt CD remains an unmet medical need^3, 4, 24, 25^. The health burden of CD in terms of quality of life, complications, mortality and cost of treatment are considerable, meaning that its prevention has become, in the last decade, an important area of research.

Metabolomics is the study of small (< 1500 Dalton) molecules and their functions in cells, tissues and body fluids^26^. Metabolomic studies in adults diagnosed with active CD^27^ identified marked changes in serum and urine metabolic profiles, along with altered intestinal microbiota^28-30^. Interestingly, a recent, small prospective study suggests an altered early trajectory of the gut microbiome (4 and 6 months of age) in children who later progressed to CD^31^. Given the characteristic metabolic abnormalities observed in clinical CD, we investigate here whether distinct lipidomic signatures exist prior to overt CD. We analyzed molecular lipids in plasma by lipidomics approach in a prospective cohort of children, observed from birth until clinical CD.

## MATERIALS AND METHODS

### Study design and protocol

The children included in the present study are from the DIPP cohort, which is an ongoing prospective study initiated in 1994. In the DIPP study, parents of newborn infants at the university hospitals of Turku, Tampere and Oulu in Finland are asked for permission to screen the child for HLA alleles conferring risk of T1D, using umbilical cord blood. Families of children identified as having an increased HLA-conferred risk for T1D are invited to join the study. Our analysis included children born at Tampere University Hospital between August 1999 and September 2005. During that period, 23,839 children were screened at birth for increased risk of T1D, and 2,642 eligible children were enrolled in the follow-up and had at least two visits to the study clinic. These children carry the high-risk HLA DQB1*02/*03:02 genotype or the moderate-risk HLA-DQB1*03:02/x genotype (x ≠ DQB1*02, 03:01, or 06:02). More than 1,200 of these children took part in the DIPP-CD study. The DIPP study in Tampere followed the children at regular intervals at the ages of 3, 6, 12, 18, and 24 months and subsequently at intervals of 12 months for children without T1D-related autoantibodies. At each visit, the families were interviewed for diet, infections, growth, important family-related issues and the children gave a non-fasting venous blood sample. The children were followed for four CD-related antibodies: anti-tissue transglutaminase (anti-tTG), anti-endomysium (EMA), antigliadin (AgA-IgG and AgA-IgA) and anti-reticulin (ARA) antibodies, and for T1D-associated autoantibodies: islet cell antibodies (ICA). If a sample was positive for ICA, then insulin autoantibodies (IAA), antibodies against tyrosine phosphatase-like protein (IA-2A) and glutamate decarboxylase (GADA) were measured from all samples taken from that child. From the beginning of 2003, all samples were measured for the four T1D-associated autoantibodies.

All children participating in this study were of Caucasian origin. IgA deficiency was excluded. One of the mothers had CD. The children in the CD follow-up cohort were annually screened for anti-tTG antibodies (tTGA) using a commercial kit (Celikey Pharmacia Diagnostics, Freiburg, Germany). If any given child’s sample was found to be positive for tTGA, all of that child’s previous and following samples were analyzed for the entire set of CD-related antibodies. A duodenal biopsy was recommended for all tTGA-positive children. If the biopsy was consistent with the ESPGHAN criteria of 1990, a gluten-free diet (GFD) was recommended. Three of the CD cases were also diagnosed with T1D, one just before the diagnosis of CD and the other ones after the diagnoses of CD (62 and 76 months later).

We randomly selected 23 children (12 males and 11 females) with biopsy-proven CD (progressors) and a control for each progressor matched for age, gender, and with the same risk HLA alleles, and living in the Tampere region throughout the whole follow-up period. None of the controls were diagnosed with T1D. The original HLA screening for CD patients and suitable controls was also completed to a “full house” DQ typing to verify the presence of DQA1*05 in selected DQB1*02 positive children^32^. These clinical and genetic annotations of the participants are given in **Table 1**. Furthermore, the children were exclusively breastfed until 1.6 months (median) and some were exclusively breastfed up to 8.0 months of age. The first exposure to gluten in this study was at a median age of 6.0 months. The children (case and control) had comparable (nominal, non-significant difference) energy (KJ), fat (g), carbohydrate (oat, rye, wheat), and gluten intakes (see **Supplementary Figures S1 & S2)**.

**Table 1.**
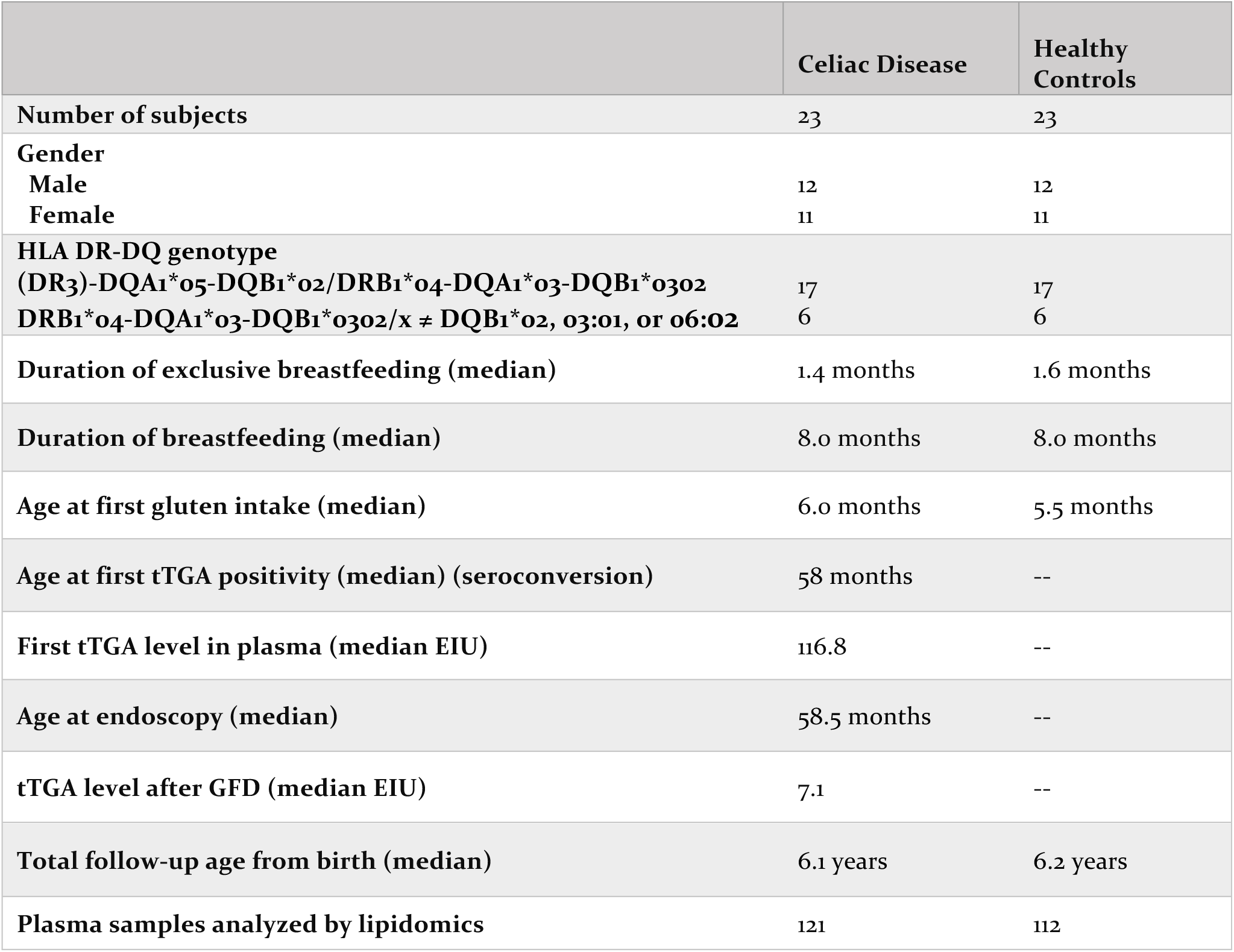
Demographic and clinical characteristics of the study subjects.

Maternal total diet during pregnancy and lactation was ascertained by a validated food frequency questionnaire^33^. Infant feeding was studied by structured questionnaires, which families filled in at home, and which were checked during the clinic visits. The child’s diet was assessed with 3-day food records at 3, 6, and 12 months followed by 2, 3, 4 and 6 years of age visits. The food records were posted before the study visit or given at the previous visit with detailed, written instructions. The families and daycare personnel were instructed to write down all the foods, drinks and dietary supplements the child consumed, with the amounts and brand names during one weekend day and two week-days. The study nurses checked the records at the study visit and helped to estimate any incomplete portion sizes with a picture booklet. The collection, processing and calculation of food consumption data has been described previously in detail^34^. The mothers of CD progressors and controls had comparable body mass indices (BMIs) and energy, fat and carbohydrate intakes during pregnancy and lactation **(Supplementary Figures S3 & S4)**.

Altogether, 121 plasma samples from children developing CD and 112 plasma samples from matched, healthy controls were analyzed, averaging more than five prospective follow-up samples for each child. Log-transformed intensities of the lipid measurements from these are given in **Supplementary Figures S5 & S6**.

### Analysis of molecular lipids

A total of 233 plasma samples were randomized and extracted using a modified version of the Folch procedure^35^. Promptly after extraction, 10 µL of 0.9% NaCl and 120 µL of CHCl_3_: MeOH (2:1, v/v) containing 2.5 µg mL^-1^ internal standard solution (for quality control (QC) and normalization purposes) were added to 10 µL of each plasma sample. The standard solution contained the following compounds: 1,2-diheptadecanoyl-*sn*-glycero-3-phosphoethanolamine(PE(17:0/17:0)),N-heptadecanoyl-D-*erythro*-sphingosylphosphorylcholine (SM(d18:1/17:0)), N-heptadecanoyl-D-*erythro*-sphingosine (Cer(d18:1/17:0)), 1,2-diheptadecanoyl-*sn*-glycero-3-phosphocholine (PC(17:0/17:0)), 1-heptadecanoyl-2-hydroxy-*sn*-glycero-3-phosphocholine (LPC(17:0)) and 1-palmitoyl-d31-2-oleoyl-*sn*-glycero-3-phosphocholine (PC(16:0/d31/18:1)), 1-palmitoyl-d31-2-oleoyl-sn-glycero-3-phosphocholine (PC (16.0/d31/18:1) that were purchased from Avanti Polar Lipids, Inc. (Alabaster, AL, USA), and tripalmitin-triheptadecanoylglycerol (TG(17:0/17:0/17:0)) (Larodan AB, Solna, Sweden). The samples were vortexed and incubated on ice for 30 min after which they were centrifuged (9400 × *g*, 3 min, 4 °C). 60 µL from the lower layer of each sample was then transferred to a glass vial with an insert, and 60 µL of CHCl_3_: MeOH (2:1, v/v) was added to each sample. The samples were re-randomized and stored at -80 °C until analysis.

Calibration curves using 1-hexadecyl-2-(9Z-octadecenoyl)-sn-glycero-3-phosphocholine (PC(16:0/18:1(9Z))), 1-(1Z-octadecenyl)-2-(9Z-octadecenoyl)-sn-glycero-3-phosphocholine (PC(16:0/16:0)), 1-octadecanoyl-sn-glycero-3-phosphocholine (LPC(18:0)), (LPC18:1), PE (16:0/18:1),(2-aminoethoxy)[(2R)-3-hydroxy-2-[(11Z)-octadec-11-enoyloxy]propoxy]phosphinic acid (LysoPE (18:1)), N-(9Z-octadecenoyl)-sphinganine (Cer (d18:0/18:1(9Z))), 1-hexadecyl-2-(9Z-octadecenoyl)-sn-glycero-3-phosphoethanolamine (PE (16:0/18:1)) (Avanti Polar Lipids, Inc.), 1-Palmitoyl-2-Hydroxy-sn-Glycero-3-Phosphatidylcholine (LPC(16:0)) and 1,2,3 trihexadecanoalglycerol (TG16:0/16:0/16:0), 1,2,3-trioctadecanoylglycerol (TG(18:0/18:0/18:0)) and ChoE (18:0), 3β-Hydroxy-5-cholestene 3-linoleate (ChoE(18:2)) (Larodan AB, Solna, Sweden), were prepared to the following concentration levels: 100, 500, 1000, 1500, 2000 and 2500 ng mL^-1^ (in CHCl_3_:MeOH, 2:1, v/v) including 1000 ng mL^-1^ of each internal standard.

The samples were analyzed using an established ultra-high-performance liquid chromatography quadrupole time-of-flight mass spectrometry method (UHPLC-QTOF-MS). The UHPLC system used in this work was a 1290 Infinity system from Agilent Technologies (Santa Clara, CA, USA). The system was equipped with a multi sampler (maintained at 10 °C), a quaternary solvent manager and a column thermostat (maintained at 50 °C). Separations were performed on an ACQUITY UPLC® BEH C18 column (2.1 mm × 100 mm, particle size 1.7 µm) by Waters (Milford, USA).

The mass spectrometer coupled to the UHPLC was a 6545 quadrupole time of flight (QTOF) from Agilent Technologies interfaced with a dual jet stream electrospray ion (dual ESI) source. All analyses were performed in positive ion mode and MassHunter B.06.01 (Agilent Technologies) was used for all data acquisition. QC was performed throughout the dataset by including blanks, pure standard samples, extracted standard samples and control plasma samples. Relative standard deviations (% RSDs) for lipid standards representing each lipid class in the control plasma samples (*n* = 8) and in the pooled plasma samples (*n* = 20) were, on average, 11.7% (raw variation). The lipid concentrations in the pooled control samples was, on average, 8.4% and 11.4% in the standard samples. This shows that the method is reliable and reproducible throughout the sample set.

MS data processing was performed using the open-source software, MZmine 2.18^36^. The following steps were applied in the processing: (1) Crop filtering with a m/z range of 350 – 1200 m/z and a RT range of 2.0 to 15.0 min, (2) Mass detection with a noise level of 1000, (3) Chromatogram builder with a min time span of 0.08 min, min height of 1200 and a m/z tolerance of 0.006 m/z or 10.0 ppm, (4) Chromatogram deconvolution using the local minimum search algorithm with a 70% chromatographic threshold, 0.05 min minimum RT range, 5% minimum relative height, 1200 minimum absolute height, a minimum ration of peak top/edge of 1.2 and a peak duration range of 0.08 -5.0, (5) Isotopic peak grouper with a m/z tolerance of 5.0 ppm, RT tolerance of 0.05 min, maximum charge of 2 and with the most intense isotope set as the representative isotope, (6) Peak list row filter keeping only peaks with a minimum of 10 peaks in a row, (7) Join aligner with a m/z tolerance of 0.009 or 10.0 ppm and a weight for of 2, a RT tolerance of 0.1 min and a weight of 1 and with no requirement of charge state or ID and no comparison of isotope pattern, (8) Peak list row filter with a minimum of 53 peak in a row (10% of the samples), (9) Gap filling using the same RT and m/z range gap filler algorithm with an m/z tolerance of 0.009 m/z or 11.0 ppm, (10) Identification of lipids using a custom database search with an m/z tolerance of 0.009 m/z or 10.0 ppm and a RT tolerance of 0.1 min, (11) Normalization using internal standards (PE (17:0/17:0), SM (d18:1/17:0), Cer (d18:1/17:0), LPC (17:0), TG (17:0/17:0/17:0) and PC (16:0/d30/18:1)) for identified lipids and closest ISTD for the unknown lipids, followed by calculation of the concentrations based on lipid-class concentration curves, (12) Imputation of missing values by half of the row’s minimum.

### Statistical methods

The samples from CD progressors and controls were divided into different age groups based on time difference between the date of the sample withdrawn and date of birth of the subject **(Supplementary Figure S7)**. If more than two samples from the same case matched a time interval, the closest was taken. The data was then log2 transformed. Homogeneity of the samples were assessed by principal component analysis (PCA)^37^ and no outliers were detected.

The R software (http://www.r-project.org) was used for data analysis and visualization. PCA was performed using *‘prcomp()’* function included in the *‘stats’* package. The effect of different factors such as age, gender and condition (healthy or CD) on the lipidomics dataset was evaluated. The data was centered to zero mean and unit variance. The relative contribution of each factor to the total variance in the dataset was estimated by fitting a linear regression model, where the normalized intensities of metabolites were regressed to the factor of interest, and thereby median marginal coefficients (R^2^) were estimated. This analysis was performed using *‘scater’* package **(Supplementary Figure S8)**. Partial least squares Discriminant Analysis (PLS-DA)^38-40^. and Variable Importance in Projection (VIP) scores^41^ were estimated by an array of functions coded in *‘ropls’* package. Moreover, the PLS-DA models were cross-validated^42^ by 7 fold cross-validation as implemented in *‘ropls’* package and Q^2^ (an estimate of model’s predictability) and R^2^ were obtained.

The longitudinal profiles of the lipids in the samples obtained from CD progressors and matched healthy controls were compared using linear mixed-effects (LME) models^43, 44^ as implemented in the *‘lme()’* function of ‘nlme’ package. The intensity of a metabolite (y) in a sample (j) is a function of multiple factors such as follow-up age, gender, case/control, subject-wise variability, etc. The clinical and genetic factors of the children were matched and standardized in this study (**Table 1**), other factors such as subject-wise variability can be modelled and derived using LME models. The LME models were restricted to constant terms with the fixed effect being CD progression/non-progression (control), age, gender and the random effect being subject-wise variation in comparison to the group-specific mean level. Fitted LME models showed, gender difference has nominal effect on the metabolite intensities. The fully parametrized model was compared to a null model using analysis of variance (ANOVA)^45^ *(‘aov()’* as deployed in the *‘stats’* package). The lipid profiles that changed significantly (p < 0.05) were subjected to False Discovery Rates (FDR) adjustment using *p-adjust()*. Lipid profiles with FDR < 0.05 were listed.

Next, post-hoc analysis using Tukey’s test for Honest Significant Difference (HSD) was performed on these selected lipids to see, if at all, they are changing between progressors and non-progressors at a particular age. A list of differentially altered lipids that also showed HSD (p-values < 0.05) between CD progressor and healthy control at a particular age was marked. These lipids were considered for further analysis.

HSD was performed using *‘TukeyHSD()’* function deployed in the ‘stats’ package. Loess regression was performed using *‘loess()’* deployed in the ‘stats’ package. Spearman’s correlation coefficient was calculated using *‘rcorr()’* function implemented in *‘Hmisc’* package. *‘Heatmap.2()’* and *‘boxplot()’* was used for data visualization.

## RESULTS

### Global plasma lipidome in progression to celiac disease

The complete lipidomics dataset was first explored using multivariate analysis. Among other factors affecting the lipidome, age was found to be a major confounding factor **(Supplementary Figure S8)**. Partial least square discriminant analysis (PLS-DA)^38-40^ of lipidomics data from all 233 longitudinal samples suggested that children who later progressed to CD (progressors) may have different lipid profiles in comparison to their matched, healthy controls (**Figure 1a)**. Different classes of lipids such as cholesterol esters (CEs), phosphatidylcholines (PCs), lysophosphatidylcholines (LPCs), phosphatidylethanolamines (PEs), phosphatidylinositols (PIs), sphingomyelins (SMs) and triacylglycerols (TGs) were affected (regression coefficient, RC (± 0.05) and VIP scores^41^ > 1) (**Figure 1b)**.

**Figure 1.**
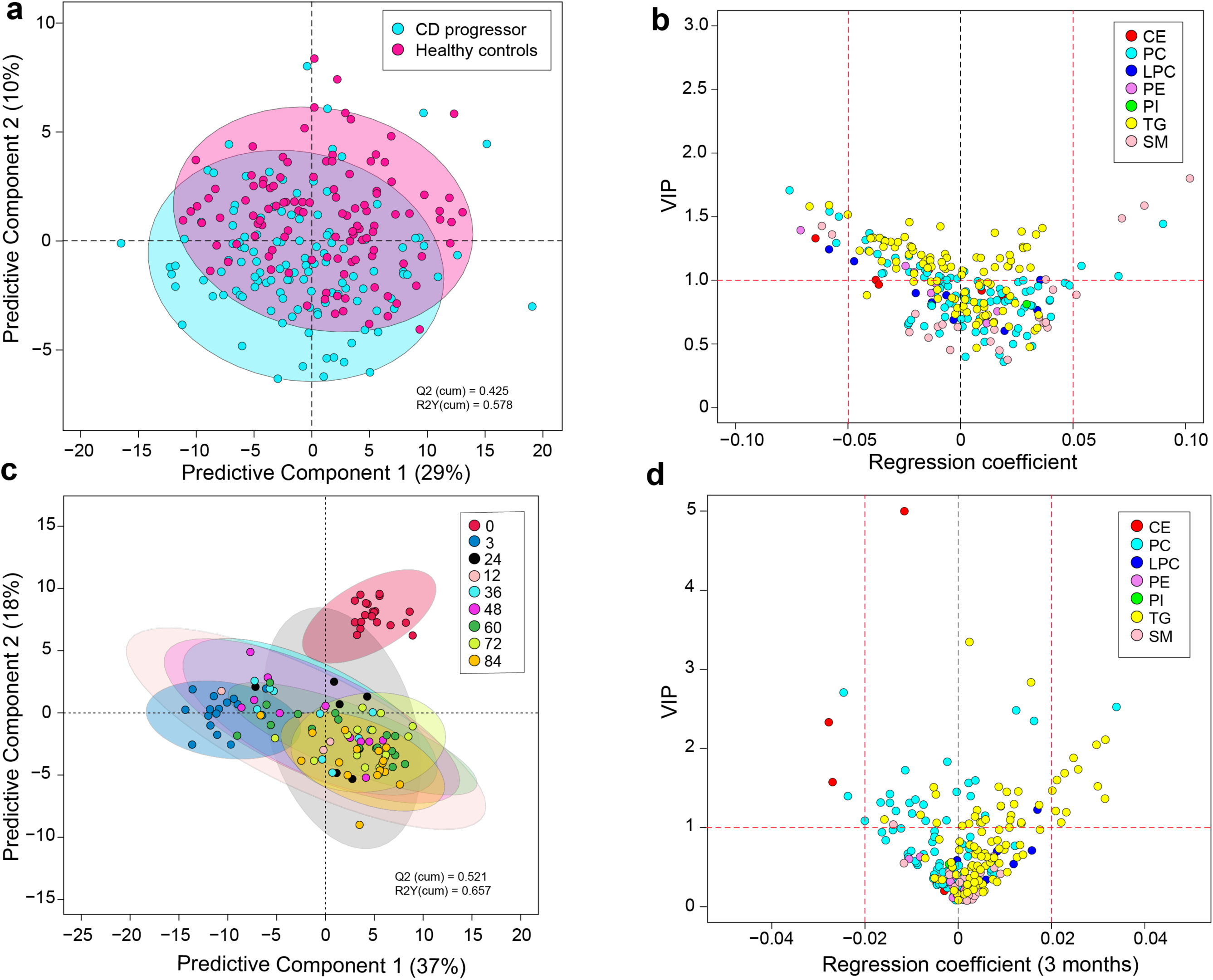
(a) PLS-DA score plot showing difference in the lipidome between CD progressors (cyan) and healthy controls (magenta). (b) A regression plot of the lipids and their VIP scores obtained from PLS-DA model. The lipids are grouped and color coded by their classes. (c) PLS-DA score plot showing difference in the lipidome in CD progressors along the age. (d) A regression plot of the lipids and their VIP scores at 3 months of age. The lipids are grouped and color coded by their classes.

Differences between cases and controls were observed already by 3 months of age, *i.e.*, before the introduction of gluten to the diet (**Figure 1c)**. TGs, PCs and CEs were mostly affected (RC (± 0.05) and VIP^41^ scores > 1) at this age (**Figure 1d)**. The cord plasma lipidome clustered distinctly from other age groups **(Supplementary Figure S9)**. However, no significant differences (HSD, p-values > 0.05) in the cord plasma lipids were observed between CD progressors and their matched healthy controls (**Figure 2a)**.

### Molecular lipids in progression to overt CD

Longitudinal analysis of lipidomic profiles at the individual lipid level identified 80 molecular lipids (from a total of 239) occurring at significantly-different levels over time between CD progressors and healthy controls (FDR-adjusted p-values < 0.05) (**Figure 2a**).

The level and direction of regulation of TGs depends on their chemical structure. There was an association between the fold change in TGs (CD progressors *vs.* healthy controls) and the TG double bond count as well as the TG carbon number (**Figure 2b-e**). At 3 months of age, TGs with lower double bond and carbon numbers were found to have increased in CD progressors as compared to matched, healthy controls, with coefficients (R= -0.13; p-values =0.19) and (R= -0.54; p-values = 0.0001) respectively for double bond count and carbon number (**Figures 2b** and **2d**). Interestingly, at a later age, these TGs with lower double bond and carbon numbers were down-regulated with (R= 0.79; p-values =0.08) and (R= 0.44; p-values = 0.0002), respectively (**Figure 2c** and **Figure 2e**). Notably, four such TGs were elevated in progressors at 3 months of age (HSD, p-values < 0.05). No significant changes were detected for dietary TGs; such as those containing polyunsaturated fatty acids (PUFAs). Representative longitudinal profiles for selected significantly-altered lipids between CD progressors and healthy controls are shown in **Figure 3**.

### Impact of tissue transglutaminase antibodies on lipidomic profiles

Tissue transglutaminase (tTG) is a multifunctional, calcium-dependent enzyme (EC 2.3.2.13) that plays an important role in the pathogenesis of CD. Antibodies produced against tTG, anti-tissue transglutaminase antibodies (tTGA), are used as serological markers with high sensitivity (99%) and reasonable specificity (> 90%) for the diagnosis of CD^2, 46, 47^.

We examined tTGA titers in 23 CD progressors at two different stages, (1) immediately after the seroconversion for tTGA, and (2) 6-12 months after both seroconversion and introduction of GFD. First, we identified at least seven ‘essential’ TGs, *i.e.*, TGs of dietary origin, in our lipidomics dataset. The identities of these TGs were validated using external sources including both the Human Metabolome Database (HMDB)^48^ and FooDB.ca (http://foodb.ca/). Spearman correlation analysis was then performed between the levels of selected TGs (34 significantly-changed, nonessential and seven non-significantly changed essential TGs) and PCs (18 PCs + 2 LPCs) with measured tTGA levels at these stages. The aim was to determine whether the systemic levels of TGs and PCs are associated with antibody titers during CD progression. A positive correlation (Spearman’s rank coefficient, δ_avg_ = +0.174; minimum p-value = 0.01) was found between the TG and tTGA levels after seroconversion (**Figure 4a**). However, a negative correlation was found at a later age, where the levels of tTGA decreased considerably (δ_avg_ = -0.163; minimum p-value = 0.05). In addition, at least 16 PCs were negatively correlated at these two different time-points (δ_avg_ = -0.02 and -0.13; minimum p-values = 0.01 and 0.02 respectively) (**Figure 4b**). It is known that GFD decreases tTGA levels in CD patients^49^, whilst it also improves the total cholesterol levels and high density lipoprotein (HDL) profiles without any significant increase in low density lipoproteins (LDL)^50^.

**Figure 4.**
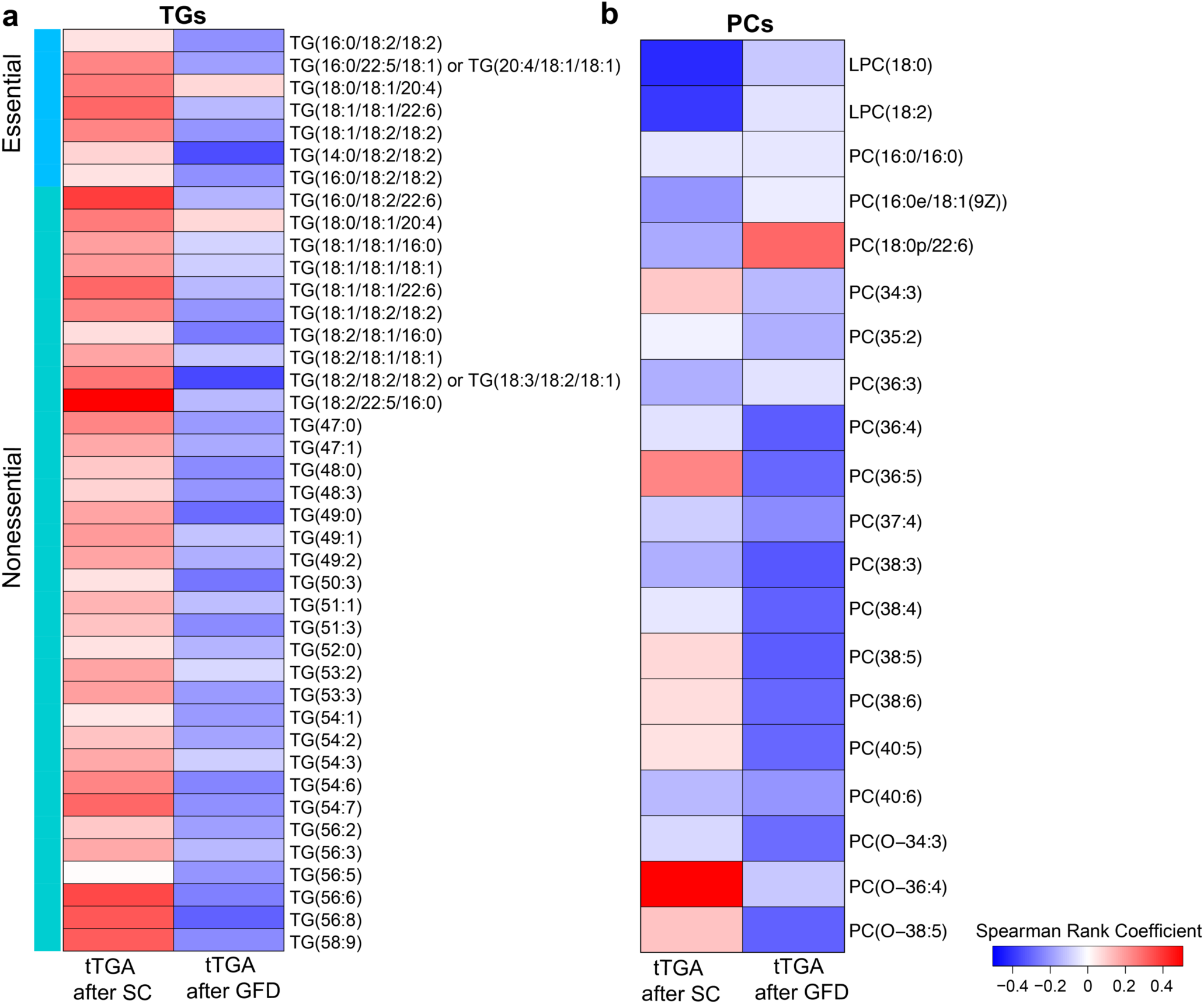
(a) Heatmap showing the Spearman’s rank coefficient estimated between significantly changed TG levels and tissue transglutaminase antibodies (tTGA) titer in the plasma of CD progressors immediately after the seroconversion (SC), and 6-12 months after the SC or introduction of gluten free diet (GFD). Red and blue color depicts positive and negative correlations. (b) Heatmap showing the Spearman’s rank coefficient estimated between significantly changed PC and plasma tTGA titer in CD progressors after SC and GFD.

### Impact of gluten on molecular lipids

We then divided the cohort into three subgroups, (1) before the introduction of gluten in the diet (3 months) (2) after the introduction of gluten (12-36 months), and (3) after the introduction of GFD (72 months, CD progressors only).

We observed a decrease in total essential TG level in the plasma of the CD progressors after gluten intake (**Figure 5a**, **Supplementary Figure S10a**). Introduction of GFD reversed this trend, but the changes were not significant (ANOVA, p-values > 0.05) (**Figure 5a**, **Figure 5e)**. On the other hand, levels of nonessential endogenous TGs were decreased (ANOVA, p-values = 0.004) in the plasma of the CD progressors after commencement of gluten intake and even after the diagnosis of clinical CD and introduction of GFD (p-values = 0.001). The trend in the change of nonessential TGs was also observed in healthy controls (p-values = 0.007 and 0.01, respectively) (**Figure 5b**, **Figure 5e** and **Supplementary Figure S10b**). Moreover, there was a difference (p-values < 0.05) in nonessential TGs levels between CD progressors and healthy controls as predicted by Tukey’s test for HSD (**Figure 2a)**. These findings suggest that dysregulation of lipid metabolism might occur at an early stage of CD progression, even before the introduction of gluten in the diet.

PCs were elevated both in the progressors (p < 0.004) and controls (p < 0.007) after the commencement of gluten intake (**Figure 5c**, **Figure 5e** and **Supplementary Figure S10c)**. PCs are the major class of phospholipids that form the cellular constituents required for the assembly of biological membranes. In contrast to healthy controls, no significant differences in SM level were observed in the CD progressors after commencement of gluten intake. A difference in the plasma SM level was observed only at a later age, after the introduction of GFD (**Figure 5d**, **Figure 5e** and **Supplementary Figure S10d**).

**Figure 5.**
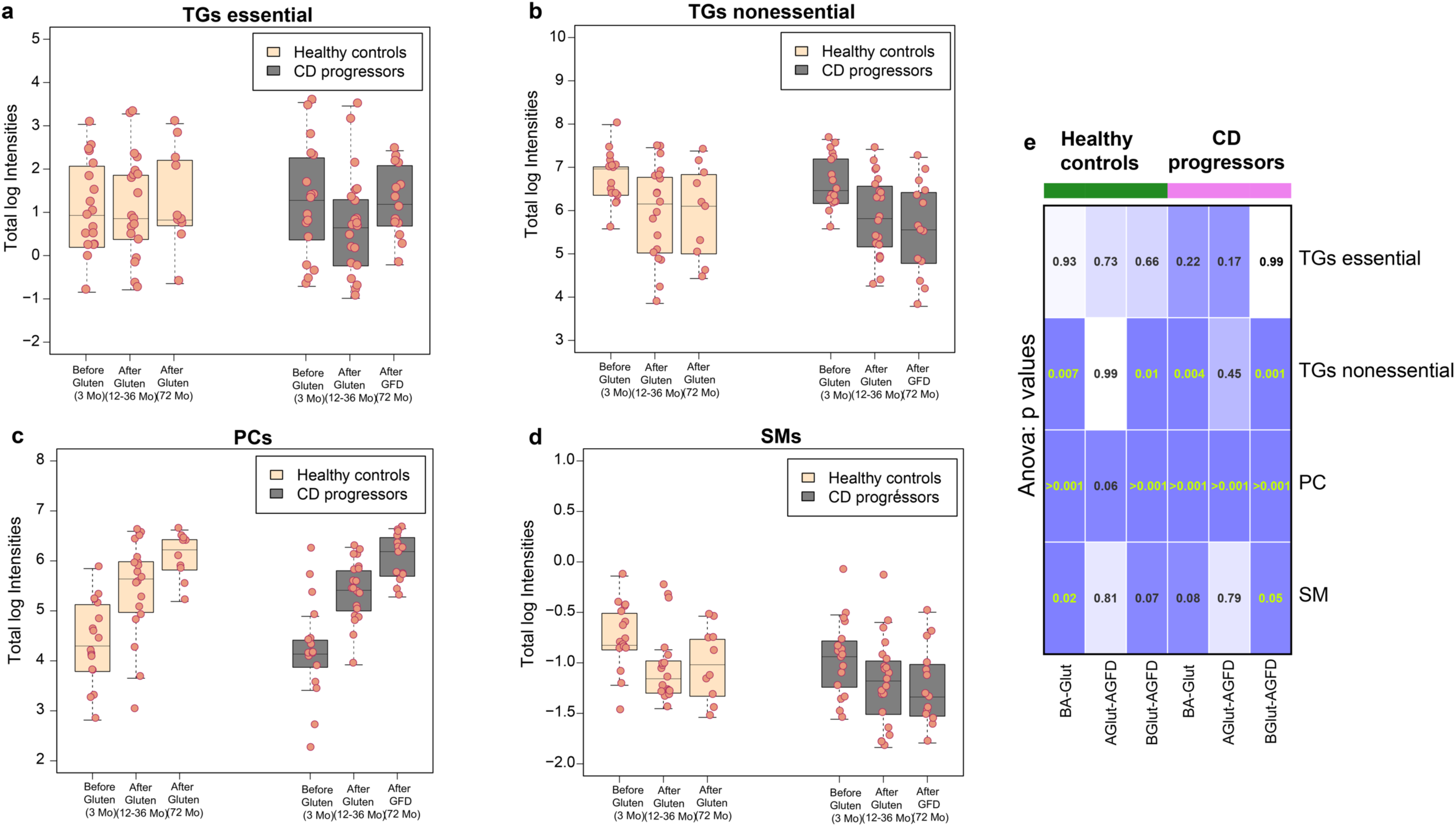
(a-d) Boxplot showing total log intensities of essential TGs, nonessential TGs, PCs and SMs in the CD progressors and healthy controls before (3 months) and after introduction of gluten (12-36 months) in the diet and after introduction of GFD (72 months). (e) Gridmap of p-values obtained from ANOVA models by combining all possible conditions before/after the gluten intake in CD progressors and healthy controls are given. The three different conditions are, *‘BA-Glut’* = before and after gluten, ‘*AGlut-AGFD’* = after gluten and after gluten free diet, *‘BGlut-AGFD’* = before gluten and after gluten free diet.

## DISCUSSION

We identified systematic differences in plasma lipidomes between children who progressed to clinical CD as compared to children who remained healthy during the follow-up. These differences were observed before the exposure to gluten in the diet and before the first signs of CD-associated autoimmunity. The dysregulation of the plasma lipidome in CD progressors is dominated by complex lipids such as PLs, TGs and CEs. As there is no evidence of gut damage in any studies performed so far in individuals who are autoantibody-negative for tTGA, it is unlikely that these systematic differences are caused by gluten from breast milk, or other sources in the infants’ diet causing damage to the gut immediately after birth. An earlier lipidomics study in the PreventCD cohort did not find significant differences between CD progressors and matched controls at 4 months of age^51^. However, that study used a targeted analysis focusing on a subset of phospholipids and acyl-carnitines, and did not measure TGs and CEs, where the major changes were found to have occurred in our study.

In this study, several nonessential TGs were up-regulated in CD progressors as early as 3 months of age, *i.e.*, before the first introduction of gluten to the diet. However, no significant changes in dietary TGs were found at this age. Lipid malabsorption is believed to be a side effect of flattened villi in the small intestine^27, 29^. However, our data suggest that lipid-related abnormalities are not caused by CD-related villous atrophy in the gut. One plausible explanation that arises is that lipid malabsorption can lead to a reduction in cholesterol-transporting lipoproteins, and secretion of apolipoprotein (Apo)-A1^52, 53^. A decrease in the level of CEs in CD progressors during the first 3 months after birth supports this explanation. Moreover, CEs were up-regulated at a later age after the introduction of GFD (**Figure 2a** and **Figure 3a**). Lipid malabsorption may thus occur at a very early age in CD progressors and, therefore, *de novo* lipogenesis^54^, as reflected by increased levels of TGs with low carbon number and double bond content, may be necessarily increased in order to compensate for said compromised lipid uptake. Interestingly, this phenomenon was not observed at birth (*i.e.*, in the cord blood data), thus suggesting that the observed lipid dysregulation is not an inborn phenomenon.

**Figure 2.**
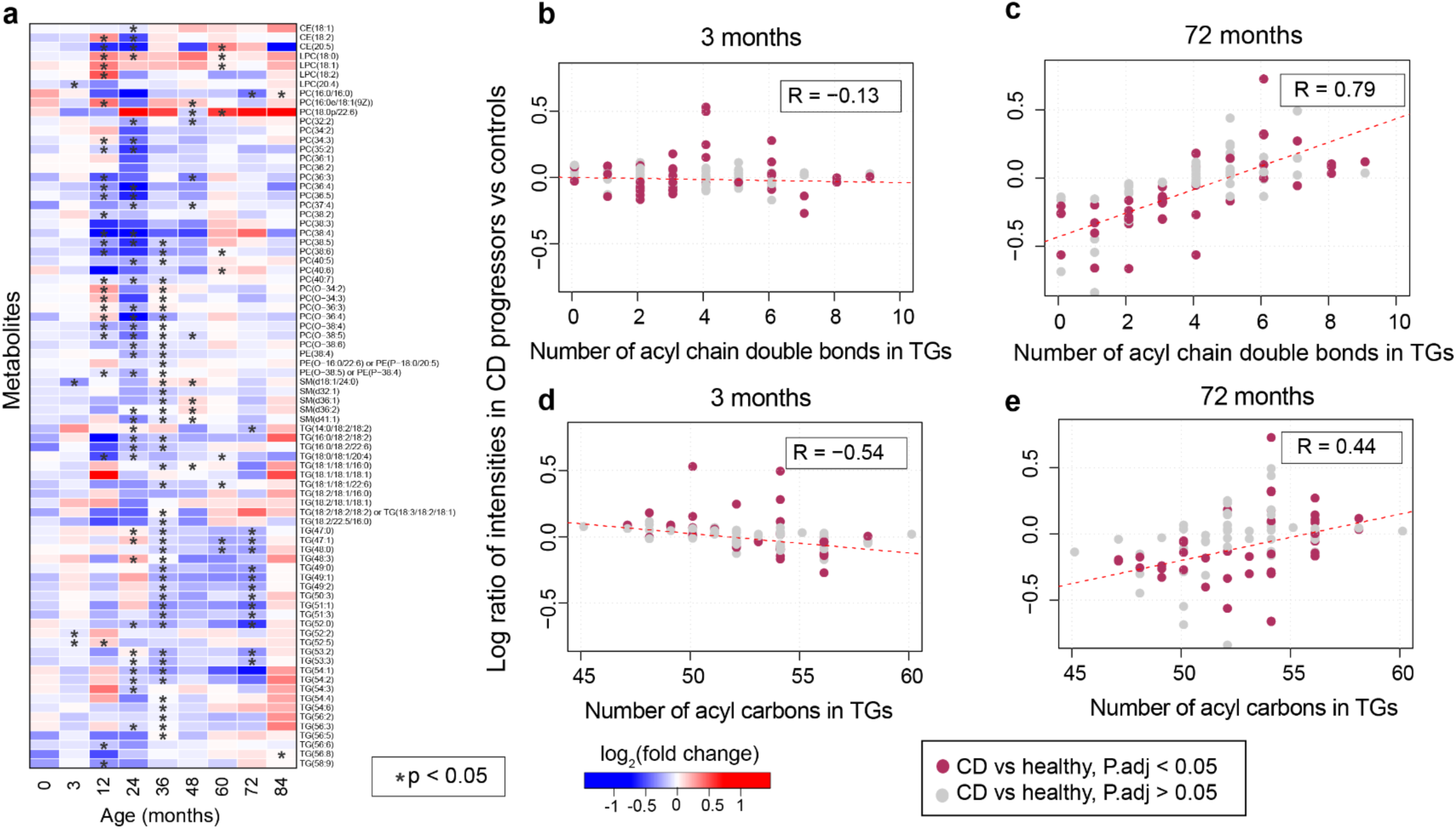
(a) Heatmap showing log_2_ fold changes (FC) of significantly altered (FDR adjusted p-values < 0.05) longitudinal lipid profiles between CD progressors and matched healthy controls, as identified by linear mixed-effect (LME) models. Here, blue and red color depicts down-and up-regulated lipid intensities in CD progressor as compared to their matched healthy controls respectively, and white depicts no change. The lipids that changed specifically (HSD, p-values < 0.05) at a particular age, as identified by Tukey’s test for HSD, is highlighted with a star ‘*’. (b-c) A correlation plot of log ration or fold changes of triacylglycerols (TGs) and the number of acyl chain bonds incurred by them at early (3 months) and later age (72 months), *i.e.*, when gluten free diet has been started. R denotes ranked correlation coefficients. (d-e) Correlation plots of log ration of TGs and the number of acyl chain carbon atoms in TGs at the same age. The significantly changed (p < 0.05) TGs between CD progressors and healthy control (at any age) are marked as dark pink color.

**Figure 3.**
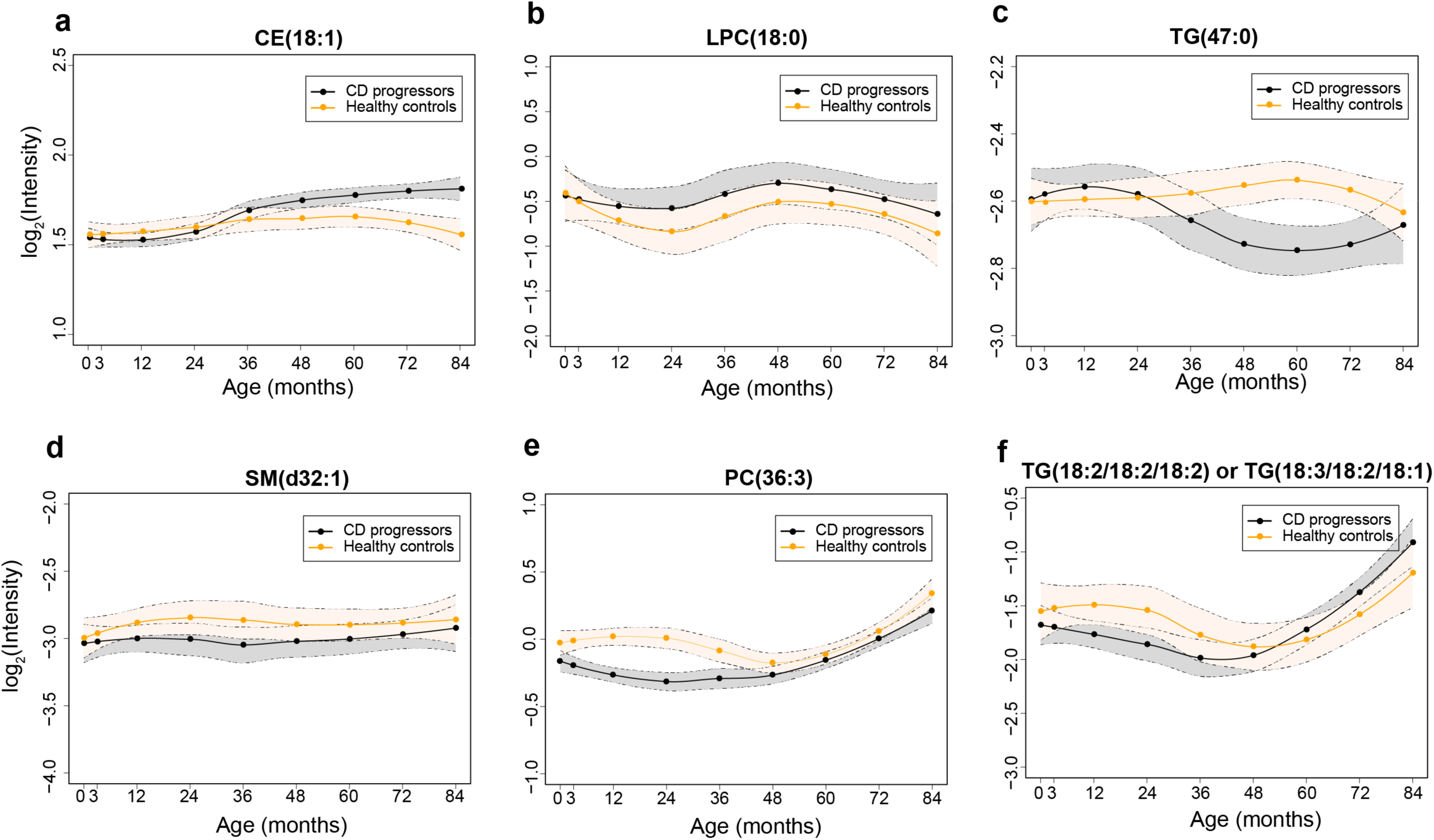
Longitudinal profiles of lipids in CD progressors (black) and healthy controls (orange) with 95% confidence interval as shown by the area shaded around the curves.

Furthermore, an inverse relationship between endogenous TGs and tTGA titer after seroconversion and introduction of GFD in the CD progressors also reaffirms the involvement of TGs and other phospholipids (PCs) in CD progression. Indeed, there was an increase in cholesterol levels after the introduction of GFD and there was a decrease in tTGA titers, in agreement with previous studies^49, 50^. Presumably, these lipids may play a protective role against seroconversion to tTGA positivity and CD progression.

We found, in earlier studies, that T1D is preceded by dysregulation of lipid metabolism^55-57^. The key findings from these studies were that islet autoimmunity and overt T1D are preceded by diminished phospholipid and TG levels. While there are some similarities at an early age in phospholipid profiles in T1D and, as shown in the current study, CD progressors, no such similarities exist for TGs. The increase of TGs with low carbon number and double bond count does appear to be specific to CD progression. These TGs are associated, in adults, with elevated liver fat in non-alcoholic fatty liver disease (NAFLD)^58, 59^, reflecting increased *de novo* lipogenesis and adipose tissue lipolysis. In fact, CD patients have, interestingly, been found to be at increased risk of NAFLD^60^. Our findings may thus offer an explanation for this association.

The main limitation of this study is the relatively small number of children included. However, this is mitigated by the longitudinal study setting, with five prospective samples, from birth until after the introduction of GFD, on average, from each child. The subjects of our study were well-matched with their controls for genetic and environmental factors.

In summary, our study suggests that lipid-related abnormalities in CD progressors demonstrably occur prior to the first introduction of gluten to the diet. These changes may be related to impaired lipid absorption as well as *de novo* lipogenesis, with both of these being part of the host response. Our findings may, therefore, have important clinical implications for the detection of subjects at-risk for CD as well as for the understanding of early pathogenesis of CD.

## DATA ACCESSIBILITY

The lipidomics datasets and the clinical metadata generated in this study have been submitted to MetaboLights^61^ and can be located using accession number (*MTBLS729*). The appropriate clinical metadata was linked to the lipidomics dataset using the ISA-creator package from MetaboLights.

## ETHICAL APPROVAL AND INFORMED CONSENT

The ethics committee of Tampere University Hospital approved the study. Written, informed consent was obtained from the parents for HLA-screening, autoantibody analysis and intestinal biopsies.

## ACKNOWLEDGEMENTS

We thank the families participating in the DIPP study for making this study possible. We also thank the expert staff of the DIPP study for their excellent work with the participating research families and sample collection. We thank Professor Olli Simell for his important scientific contribution to the DIPP study. We thank to Dawei Geng for technical assistance in lipidomic analysis and to Aidan McGlinchey for editing the manuscript.

